# A claustro-cortical loop times state transitions for flexible behavior

**DOI:** 10.64898/2026.07.10.737800

**Authors:** Randall J. Olson, Alex Sonneborn, Russell Milton, Lowell Barlett, Atheir I. Abbas

**Author notes:** Corresponding Author;* Correspondence and requests for materials should be addressed to Atheir I. Abbas.

## Abstract

Metastable dynamics, in which neural activity moves between transiently stable patterns, have been proposed to underlie the real-time coordination on which the brain’s cognitive and behavioral functions depend^1–3^. In the cortex, the simplest form of this regime is the alternation between active ‘UP’ states, in which neurons fire, and silent ‘DOWN’ states, in which firing is strongly reduced, a bistable architecture that shapes how afferent information is processed ^4–8^. Such metastable dynamics are ubiquitous across cortex ^9,10^, yet how their state transitions are timed and controlled, and how this timing shapes behavior, remain open questions. Here we show, in freely behaving mice performing a cue-guided switching task, that the metastable dynamics of frontal cortex are jointly shaped by its reciprocal loop with the claustrum, a small and widely connected subcortical structure, and that the loop’s timing of these state transitions is required for efficient flexible behavior. Claustrum activity led the cortical UP-to-DOWN state transition, delivering to anterior cingulate cortex a low-dimensional drive aligned with the axis along which the cortical state switches. This drive was carried not by a rise in claustral firing but by transient coordination: population synchrony peaked at the transition even as mean rate fell. Silencing ACC-projecting claustrum terminals reduced directed claustro-cortical coupling, biased cortex away from the DOWN state by prolonging UP states and shortening DOWN states, and selectively impaired cue-driven switching while sparing exploratory foraging. These results place flexible behavior under the control of a defined cortico-subcortical loop that times cortical state transitions, offering a circuit-level entry point into the cognitive rigidity of many psychiatric and neurological conditions^11–13^.

## Main

During waking, cortical activity is not stationary, instead fluctuating rapidly through a repertoire of discrete internal states that shape how incoming stimuli are processed ^9,10^. The consequence is substantial: the same stimulus can evoke markedly different behavioral responses depending on the state the network occupies when it arrives, a dependence documented across species, brain regions and sensory modalities^6,14,15^. This repertoire has been characterized chiefly in sensory cortex, where wakefulness resolves into a graded family of substates rather than a single mode ^10,16,16^. Its simplest and most thoroughly studied instantiation is the binary alternation between active states of sustained, synaptically driven firing (UP) and relatively quiescent states of strongly reduced firing (DOWN) ^4,5,17,18^, and we take this two-state description as our working reduction of cortical state. These two states are not points on a smooth continuum but discrete configurations: the network settles into one, persists there for a period, and then switches to the other comparatively abruptly rather than drifting between them. Dynamics of this kind, in which activity occupies a transiently stable pattern before being abandoned for the next, are termed metastable, and the UP/DOWN alternation is their simplest instance ^1,2,17,19^. Yet what generates these states and what sets the timing of the transitions between them have remained largely unresolved, and whether this state structure is actively controlled to serve behavior is essentially unknown, particularly in the prefrontal regions that guide flexible action. We therefore asked whether a defined subcortical circuit regulates transitions between these states during behavior, promoting the DOWN states that punctuate prefrontal activity and separate one active epoch from the next.

Flexible behavior depends, in part, on the anterior cingulate cortex (ACC). Lesion or inactivation of ACC selectively impairs the updating of behavioral strategy without affecting execution of learned responses ^20–22^, and individual ACC neurons track outcome history, action value and decision uncertainty, the variables used to decide when an established response should be revised ^23,24^. Yet the output of ACC is not a fixed function of these variables: under matched conditions an established response is revised on some occasions and maintained on others ^25,26^, So part of what determines whether behavior is updated may lie not in the decision variables alone but in the rapidly fluctuating repertoire of internal states that the cortex cycles through, described above^9,10,14^. UP and DOWN states are a known source of such variation, since cortical responsiveness to identical input differs sharply between them ^7,27^. What organizes these states in ACC, and what controls the timing of the transitions between them, therefore bears directly on how ACC, or any cortical region, supports flexible behavior.

These questions invite a dynamical-systems approach to cortical state. Cortical computation can be framed as motion through a landscape of metastable attractors, in which the network dwells in a transiently stable pattern of population activity for hundreds of milliseconds before transitioning, relatively abruptly, to the next ^1,2,19^; in this framing, the UP/DOWN alternation is a bistable system in which a sustained active state and a quiescent silent state each act as an attractor ^28^. A central and unresolved problem in this account is the origin of the transitions themselves: in a network whose attractors are stable by construction, what drives escape from one state to the next, and how is its timing controlled? Recent theory shows that reliable yet flexibly timed transitions arise when a small, recurrence-poor structure reciprocally coupled to the cortical network injects low-dimensional, correlated variability aligned with the transition axis in population-activity space that separates successive attractors ^29^. In this account the metastable dynamics can arise from the coupled cortico-subcortical circuit rather than of cortex alone, since the transitions are regulated within the loop rather than imposed on cortex from outside it. The motif predicts the involvement of a subcortical partner but leaves its identity open, with thalamic and basal ganglia circuits proposed as candidates. We reasoned that the claustrum is positioned to be that partner for ACC.

Several features of claustrum anatomy and physiology fit this role. It is small and lacks the recurrent excitatory connectivity that builds cortical attractors; it maintains dense reciprocal connections with neocortex and projects most heavily to prefrontal and cingulate regions^30,31^; its individual neurons integrate convergent input from multiple frontal areas, providing a substrate for reading ongoing cortical dynamics across a wide territory ^32^; and its projections back to cortex preferentially recruit interneurons ^33,34^, positioning claustrum output to deliver the correlated, low-dimensional input that could promote a transition into a DOWN state. Optogenetic activation of the claustrum drives widespread cortical DOWN states ^34^, and ACC-projecting claustrum neurons support attentional engagement and behavioral flexibility ^35–38^.Together these features suggest that the claustrum is not merely modulated by cortical activity but is positioned to act on it, delivering the kind of targeted, synchronizing input that could help drive ACC from one state to the next.

We hypothesized that state transitions within the metastable dynamics of ACC are regulated by its reciprocal circuit with the claustrum, with the claustrum timing each transition into the silent state, and that this timing is required for flexible behavior. To test this, we combined cell-type-specific photoinhibition of ACC-projecting claustrum terminals with simultaneous field-potential and single-unit recording in ACC and claustrum during a cue-guided switching task. We find that claustrum activity leads the UP-to-DOWN transition and delivers a low-dimensional drive aligned with the axis along which cortex switches states, carried not by the level of claustral firing but by a transient synchronization. Silencing this input disrupts termination of the active state, so that cortex remains abnormally locked in the UP state and optimal cue-guided switching is impaired. These results place the timing of cortical state transitions under the control of a specific subcortical partner and make that timing a requirement for optimal flexible behavior.

### Claustrum input to ACC is required for efficient cue-guided switching

Water-restricted mice were trained on a cue-guided switching task in which on each trial they foraged for small water rewards across a foraging zone (5 front ports) while a brief auditory cue, which played unpredictably across trials, signaled large-reward availability at a back port (Fig. 1a). Across training, back-port visits became selectively locked to the cue (Fig. 1b) and discrimination performance reached a median d′ of 1.60 at test (signed-rank versus 0, P = 1.3 × 10^−□^ Fig. 1c). Hit rates, defined as cued back-port visits, substantially exceeded false-alarm rates, defined as uncued back-port visits (60% versus 16%; P = 4.0 × 10^−□^, n = 19 mice; Fig. 1d), confirming that back-port visits were cue driven rather than a generalized response bias.

**Figure 1.**
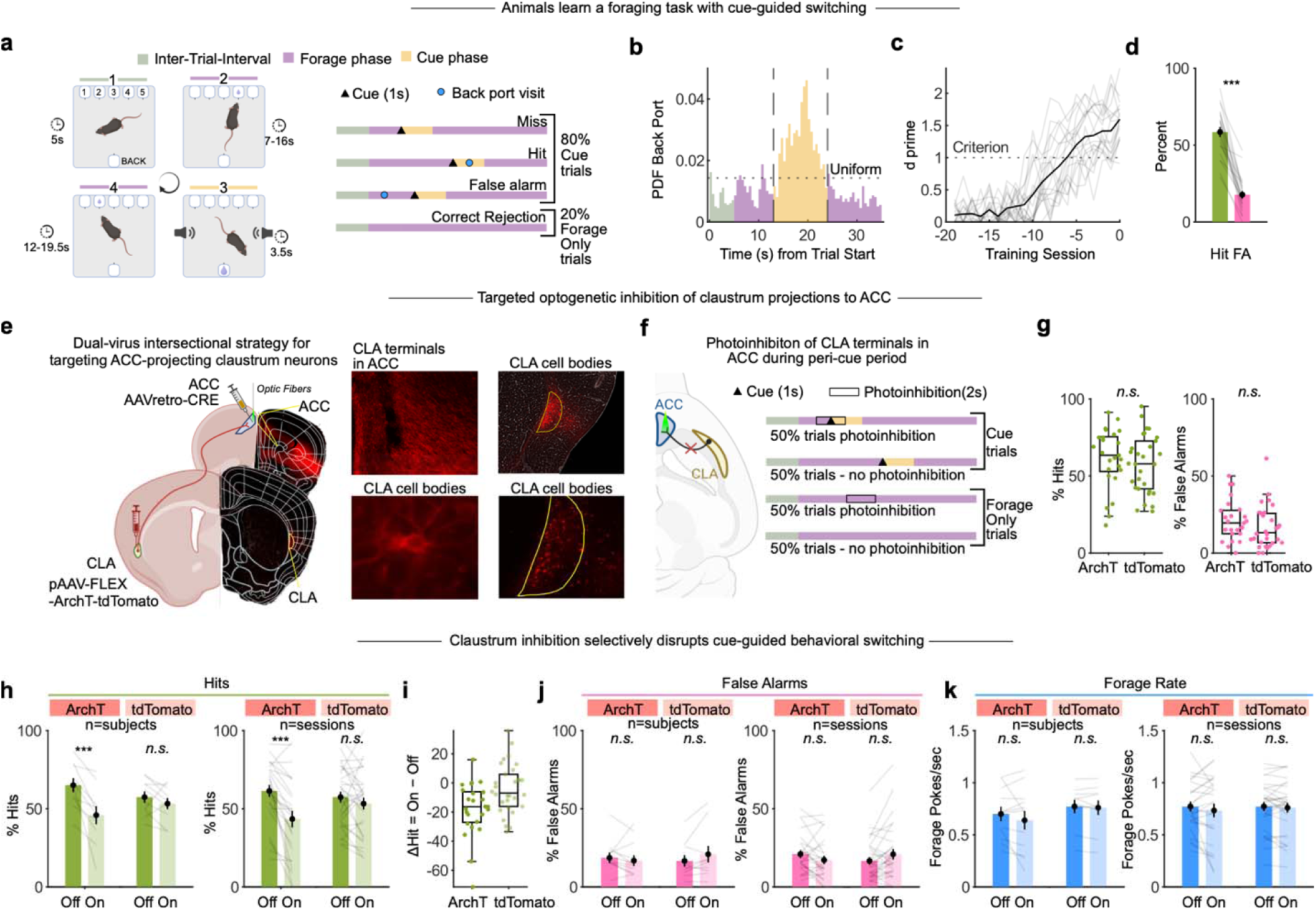
Claustrum input to anterior cingulate cortex is required for optimal cue-guided switching. (**a**) Auditory cue-guided foraging task. Water-restricted mice foraged for small rewards across a foraging zone while a brief chirp signaled large-reward availability at a back port. Mice must selectively use the cue to guide back-port selection. (**b**) Across training, back-port visits became selectively locked to the cue (n= 19 mice). (**c**) Discrimination performance at test, median d′ = 1.60 (Wilcoxon signed-rank versus 0, P = 1.3 × 10^−^□; n = 19 mice). Individual animals in light grey; black trace, average over animals. (**d**) Cued back-port visits (hits) exceeded uncued visits (false alarms), 60% versus 16% (P = 4.0 × 10^−^□, n = 19 mice), confirming cue-driven rather than biased responding. (**e**) Intersectional dual-virus strategy expressing the inhibitory opsin ArchT, or fluorophore only in controls, in ACC-projecting claustrum neurons, with bilateral optic fibers implanted over ACC for terminal photoinhibition. Inset: representative histology confirming fiber placement and terminal expression. (**f**) Optogenetic inhibition protocol. Laser was time-locked to the peri-cue period (–0.5 to +0.5 s relative to cue onset/offset; 2 s total), delivered pseudorandomly on cue and forage-only trials. (**g**) Baseline hit and false-alarm rates on laser-off trials in ArchT and control animals. No significant between-group differences (hit P = 0.393; false alarm P = 0.163). (**h**) Peri-cue photoinhibition reduced hit rate in ArchT animals (animal level 66% to 48%, P = 4.9 × 10^−^□ ; session level 63% to 45%, P = 8.1 × 10^−^□, n = 25 sessions) but not in controls (P > 0.15). (**i**) The within-session hit-rate decrement (laser-on minus laser-off) exceeded that in controls (median −16% versus −7%; between-group P = 0.0084). (**j**, **k**) False-alarm and front-port visit rates were unchanged in both groups animal and session level. Box plots show median and interquartile range.

To ask whether claustrum input to ACC is required for this behavior, we restricted the inhibitory opsin ArchT to ACC-projecting claustrum neurons using a dual-virus intersectional strategy^33^. A retrograde Cre virus (AAVrg-Cre) was injected into ACC, and a Cre-dependent ArchT virus (AAV-FLEX-ArchT-tdTomato), or a Cre-dependent tdTomato fluorophore alone in controls was injected into the claustrum, so that opsin expression was confined to claustrum neurons projecting to ACC. We then implanted optic fibers over ACC to photoinhibit their terminals selectively, leaving claustrum somata and their projections to other targets unperturbed. (Fig. 1e). Trials were of two types, cued (80%) and forage-only (20%), and laser was delivered on half the trials of each type, time-locked to the peri-cue period (Fig. 1f). Baseline performance on laser-off trials did not differ between ArchT and control animals (Fig. 1g). Peri-cue photoinhibition selectively reduced hit rates in ArchT animals (animal level 66% to 48%, P = 4.9 × 10^−□^ ; session level 63% to 45%, P = 8.1 × 10^−□^, n = 25 sessions; Fig. 1h) but had no effect in controls (P > 0.15), and the within-session decrement exceeded that in controls (median change in hit rate, −16% versus −7%; between-group P = 0.0084; Fig. 1i). False-alarm and front-port visit rates were unchanged in both groups (Fig. 1j,k), showing that photoinhibition selectively disrupted cue-guided switching without altering exploratory behavior or producing general response suppression.

### The architecture of UP/DOWN cycling tracks behavioral engagement

We next characterized the ongoing UP/DOWN state dynamics of ACC during the task. The dominant periodic component of the ACC field potential during performance was the slow oscillation in the delta band, consistent with prominent UP/DOWN cycling (Extended Data Fig.1). To analyze this fast architecture directly, we classified UP and DOWN states from concurrent LFP and multi-unit activity using a validated combined-evidence approach^39^: the analytic phase of the low-frequency ACC LFP and a smoothed multi-unit signal were combined into a single state-evidence variable, fit with a two-component Gaussian mixture model and assigned to UP or DOWN states by maximum a posteriori probability (Fig. 2a-b; Methods). This combined-evidence classification tracked UP/DOWN structure alongside behavior on individual trials (Fig. 2c). Within a single trial the architecture was not stationary: during a disengaged epoch lacking forage poke events, the network cycled rapidly and regularly between UP and DOWN states at low population rate (Fig. 2d), whereas during an engaged foraging epoch the active state became sustained and DOWN intrusions grew sparse (Fig. 2e). We next asked whether this qualitative shift held across the population.

**Figure 2.**
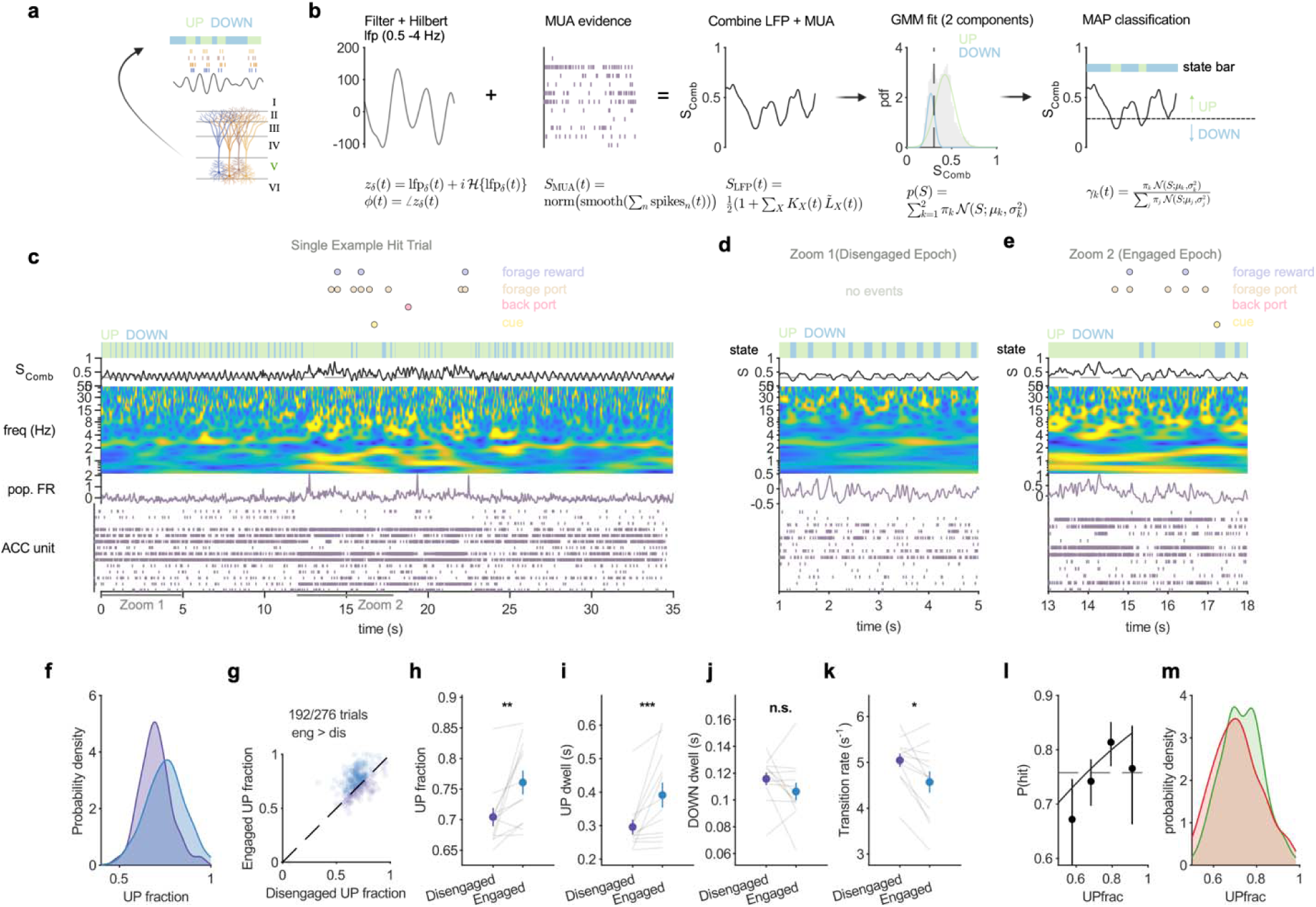
The architecture of UP/DOWN cycling tracks behavioral engagement. (**a**) Schematic of recording in deep layers of the ACC during the foraging task, with cortical activity segmented into UP (green) and DOWN (blue) states. (**b**) Classification of cortical UP and DOWN states from concurrent ACC local field potential and multi-unit activity using a combined-evidence approach (Saleem et al., 2010): the analytic phase of the low-frequency LFP (1–4 Hz) and a smoothed multi-unit signal were combined into a single state-evidence variable, fit with a two-component Gaussian mixture model and assigned by maximum a posteriori probability. (**c**) Full 35-s example trial (session 1, trial 10, hit, laser-off). From top: event raster, with forage reward (purple), forage poke (peach), cue (yellow) and cued back-port visit (pink); state bar, UP (green) and DOWN (blue); combined-evidence variable S_comb with the UP/DOWN classification threshold (dotted line); spectrogram (0.5–55 Hz, log scale); ACC population firing rate; and ACC single-unit raster, each row a neuron. (**d, e**) Same panel structure as (c), restricted to a disengaged epoch (d, 1–5 s of the same trial) and an active foraging epoch (e, 13–18 s of the same trial). (**f–k**) UP/DOWN dynamics compared between disengaged periods and active foraging within the same laser-off trials (n = 12 sessions with sufficient simultaneously recorded ACC units). (**f**) Probability density of UP fraction during foraging and disengaged epochs (Kolmogorov-Smirnov D = 0.2971, P = 3.064 × 10^-11^). (**g**) Within-trial comparison of UP fraction; foraging exceeded disengaged in 192 of 276 paired trials. (**h**) The cortex spent a greater fraction of time in the UP state during foraging (0.76 versus 0.70; signed-rank P = 0.002). (**i**) UP state dwell time was longer during foraging than during disengaged epochs (391 versus 296 ms; signed-rank P = 0.001). (**j**) DOWN state dwell time was unchanged between foraging and disengaged epochs (106 versus 116 ms; signed-rank P = 0.20). (**k**) The cortex cycled more slowly between UP and DOWN states during foraging (4.6 versus 5.0 transitions s^−1^; signed-rank P = 0.016). (**l**) Across cued trials, the probability of a correct response rose with pre-cue UP fraction and saturated (mixed-effects logistic regression, odds ratio 1.233 per s.d., P = 0.0067; Spearman ρ = 0.098, P = 0.0022). (**m**) Probability density of pre-cue UP fraction for correct versus incorrect cued trials (Kolmogorov-Smirnov D = 0.12, P = 0.0071).

We compared UP/DOWN dynamics between disengaged periods, when the animal was not sampling ports, and bouts of active foraging within the same laser-off trials (Hit and Miss pooled; n = 12 sessions with sufficient (>5) simultaneously recorded ACC units (Methods). The cortex occupied the UP state for a greater fraction of the time during foraging than during disengagement, a difference evident across the full distribution of UP fractions (Kolmogorov–Smirnov D = 0.30, P = 3.1 × 10⁻¹¹; Fig. 2f), present within trials, where foraging exceeded disengagement in 192 of 276 paired trials (Fig. 2g), and significant in the session-wise summary (0.76 versus 0.70; signed-rank P = 0.002; Fig. 2h). This redistribution reflected a lengthening of the UP state rather than a change in the DOWN state, since UP dwell times were longer during foraging (391 versus 296 ms; signed-rank P = 0.001; Fig. 2i) while DOWN dwell times were indistinguishable between conditions (106 versus 116 ms; signed-rank P = 0.20; Fig. 2j). Consistent with a selective extension of the UP state, the cortex cycled more slowly between UP and DOWN states during foraging (4.6 versus 5.0 transitions s^−1^; signed-rank P = 0.016; Fig. 2k), so that engagement was accompanied by a shift toward more sustained cortical activation rather than faster state alternation. This architecture was related to behavior on a trial-by-trial basis, since across cued trials the probability of a correct response rose with the fraction of the pre-cue window that the cortex had spent in the UP state and then saturated (mixed-effects logistic regression, odds ratio 1.233 per s.d., P = 0.0067; Spearman ρ = 0.098, P = 0.0022; Fig. 2l), indicating that the state the cortex occupied before the cue was predictive of the animal’s subsequent choice, a relationship also apparent in the distribution of pre-cue UP fraction, which was shifted toward higher values on correct than on incorrect trials (Kolmogorov-Smirnov D = 0.12, P = 0.0071; Fig. 2m). Together these results show that the UP/DOWN architecture of ACC is not fixed but tracks behavioral engagement and predicts performance. This raised the question of what controls the timing of transitions into the DOWN state during engaged behavior.

### Claustrum delivers a low-dimensional, transition-aligned drive that leads the cortical UP-to-DOWN transition

To ask whether the claustrum supplies that drive, we recorded claustrum and ACC simultaneously and examined claustral activity around individual cortical UP-to-DOWN transitions. Using the same UP/DOWN classification as above (Fig. 2; Methods), we aligned spiking in each area to every cortical transition, restricting all analyses to the active-foraging epochs in which cue-guided switching occurs and confining each estimate to the bounds of the departing and arriving states, so that no estimate could draw on a neighboring cycle. To display the population without imposing structure, we ordered neurons by a cross-validated estimate of peak latency, sorting on one half of each neuron’s transitions and showing the held-out half. Aligned to their own transitions, ACC units showed the expected step in z-scored rate, sustained through the UP state and dropping sharply at the UP-to-DOWN transition, with the mirror-image rise at the DOWN-to-UP transition (Fig. 3a). Aligned to the same cortical transitions, the claustral population showed a coherent but smoother and lower-amplitude modulation, predominantly UP-active and falling across the UP-to-DOWN transition, with no ordered single-neuron sequence (Fig. 3b). Claustral firing thus tracked the cortical state as a distributed, low-dimensional population signal.

**Figure 3.**
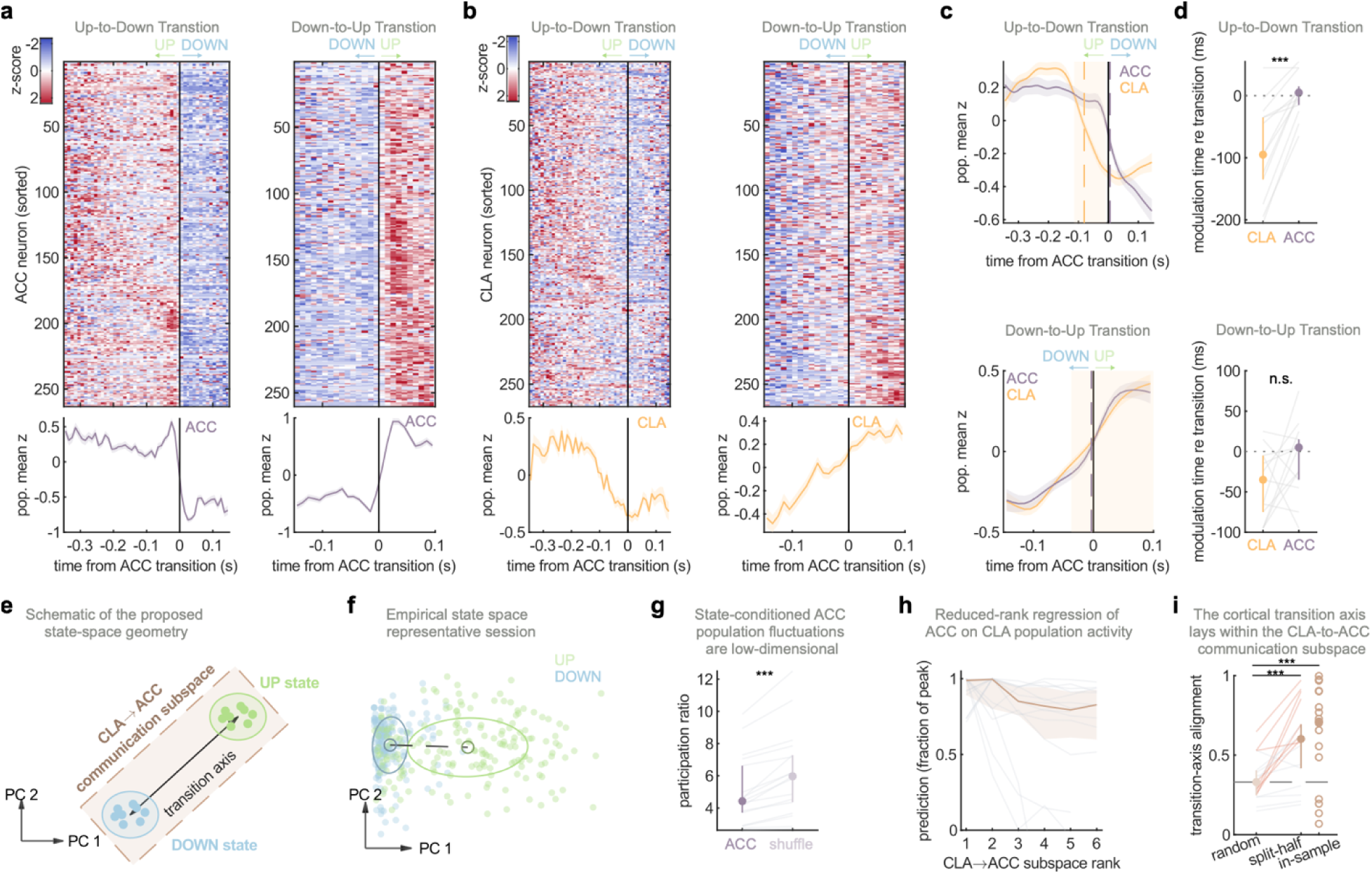
Claustrum activity leads the cortical UP-to-DOWN transition and supplies a low-dimensional, axis-aligned drive. (**a**) Cross-validated population heatmap of ACC single-unit activity aligned to ACC UP-to-DOWN (left) and DOWN-to-UP (right) transitions; neurons are ordered by their response in a held-out half of transitions and displayed on the complementary half, so the ordering imposes no apparent structure. Beneath each heatmap, the population-mean z-scored trace across ACC neurons. (**b**) Cross-validated population heatmap of CLA single-unit activity aligned to the same UP-to-DOWN (left) and DOWN-to-UP (right) transitions, ordered and displayed as in (a). Beneath each heatmap, the population-mean z-scored trace across CLA neurons. (**c**) Population-mean traces from (a) and (b) overlaid (ACC, purple; CLA, orange) at the UP-to-DOWN (top) and DOWN-to-UP (bottom) transitions. Cross-correlation of the two traces yields a claustrum lead of 86 ms at the UP-to-DOWN transition (positive values indicate claustrum leading), which persisted when cortical states were detected from a held-out subset of ACC units, excluding shared-unit circularity (session-clustered bootstrap 95% CI 14 to 120 ms, P = 0.016, n = 15 sessions). No resolvable lead was present at the DOWN-to-UP transition (P = 0.48). (**d**) Per-session comparison of the time of steepest change in the population-mean CLA and ACC traces relative to each transition, with transitions detected from a held-out subset of ACC units so that neither estimate is fixed at the transition by construction. At the UP-to-DOWN transition (top) the claustrum trace turned before that of ACC in 12 of 13 sessions (claustrum −95 ms versus ACC 5 ms; median lead 90 ms; signed-rank P = 0.0005), consistent with the cross-correlation in (c). At the DOWN-to-UP transition (bottom) the claustrum did not reliably precede ACC (claustrum −35 ms versus ACC 5 ms; median lead 20 ms; signed-rank P = 0.146), so the lead is specific to the UP-to-DOWN transition. (**e**) Schematic of the proposed state-space geometry. ACC population activity is represented in a low-dimensional state space spanned by its leading principal components, within which the UP and DOWN states occupy distinct regions and the change between them defines a single direction, the transition axis. The CLA-to-ACC communication subspace, the low-dimensional set of ACC dimensions predicted by claustral activity, is the shaded region containing the transition axis. Panels (g) to (i) test the three elements of this scheme in turn: the dimensionality of the ACC fluctuations, the rank of the communication subspace, and the alignment of the transition axis with it. (**f**) Empirical state space from a representative session. ACC population activity projected onto its leading principal components, each point colored by the concurrent cortical state (UP, green; DOWN, blue). The states separate along a single direction, the empirical counterpart of the transition axis drawn in (e). (**g**) State-conditioned ACC population fluctuations were low-dimensional: the participation ratio of the noise covariance was below that of a covariance-destroying shuffle (4.43 versus 5.97; signed-rank P = 2.9 × 10^−^□ ; n = 17 sessions). (**h**) Reduced-rank regression of ACC on CLA population activity identified a low-rank communication subspace, with prediction saturating within two dimensions (median optimal rank 2). (**i**) The cortical transition axis was aligned the CLA-to-ACC communication subspace above a random-subspace null (in-sample alignment 0.71 versus 0.33, P = 1.2 × 10^−3^), and remained so under a split-half estimate taking the axis and the subspace from independent epochs (0.60 versus 0.33, P = 1.0 × 10^−3^; above chance in 8 of 17 sessions individually).

The claustral population mean, however, appeared to turn before the cortical one (Fig. 3c,Top). To test this while avoiding circularity, we noted that the ACC multiunit signal contributes to state detection, so a direct comparison would be confounded; we therefore detected states from a randomly selected half of the ACC units and aligned the complementary, held-out half to the resulting transitions, so that neither the claustral units nor the compared cortical units informed the placement of the transition. Under this procedure the claustral population led the cortical UP-to-DOWN transition by 86 ms (cross-correlation of population-mean traces; session-clustered bootstrap 95% CI 14 to 120 ms; one-sided P = 0.016, n = 15 sessions; Fig. 3c,Top). An independent session-wise estimate agreed: the time of steepest change in the claustral trace preceded that of ACC in 12 of 13 sessions (median lead 90 ms; signed-rank P = 0.0005; Fig. 3d,Top). The DOWN-to-UP transition showed no resolvable claustral lead by either method (cross-correlation P = 0.48; session-wise median 20 ms, P = 0.146; Fig. 3c,d,Bottom), an asymmetry to which we return below.

These results identify the claustrum as a candidate directed source of a transition drive but do not characterize its geometry within ACC population activity. The metastable-attractor framework makes a specific prediction: the variability driving transitions should be low-dimensional, and its principal direction in population-activity space should align with the axis separating the states it connects (Recanatesi et al., 2022; Fig. 3e). We tested this in the simultaneously recorded populations (n = 17 sessions with at least five simultaneous units per area). Within a session, ACC population activity separated into UP and DOWN regions along a single direction, the transition axis (Fig. 3f). State-conditioned ACC fluctuations were low-dimensional: the participation ratio of the noise covariance was well below that of a covariance-destroying shuffle (4.43 versus 5.97; signed-rank P = 2.9 × 10^−^□; Fig. 3g). Reduced-rank regression of ACC on claustrum activity identified a communication subspace of low cross-validated rank, with prediction saturating within two dimensions (median optimal rank 2; Fig. 3h), confirming that the shared claustrum-to-ACC variability is low-dimensional. Finally, this drive aligned with the transition axis: the axis lay substantially within the claustrum-to-ACC communication subspace, far more than expected for a random subspace of equal dimension (in-sample alignment 0.71 versus null 0.33; signed-rank P = 1.2 × 10^-3^; Fig. 3i). Because estimating the axis and the subspace from the same epochs could inflate this, we repeated it split-half, taking the axis from one set of epochs and the subspace from another; alignment remained well above null (0.60 versus 0.33; P = 1.0 × 10^-3^), exceeding chance in 8 of 17 individual sessions. Together, these data support the claustrum as a transition-leading input to ACC that is low-dimensional and oriented, in population space, along the axis the cortex travels to reach the DOWN state.

### A coordinated claustral volley, not a rise in firing rate, accompanies the UP-to-DOWN transition

Having shown that the claustrum leads the cortical UP-to-DOWN transition and is oriented along the transition axis (Fig. 3), we asked what feature of claustral activity carries that drive. The simplest explanation, that claustral firing ramps up to push cortex into the DOWN state, was not met: the population mean rate fell as the cortex approached the DOWN state, declining from its UP-state level into the transition and reaching its minimum at the transition itself (Fig. 3c). The drive that leads the transition is therefore not an increase in mean firing.

We reasoned that the drive might instead be carried by the coordination of claustral firing rather than its mean level, since the mean rate and the population synchrony of an ensemble are distinct quantities that can move independently: a brief, coordinated population event can occur even as the average rate falls (Fig. 4a). To isolate coincident firing beyond that expected from the rates alone, we computed cross-neuron spike coincidence in each peri-transition bin and subtracted a rate-preserving null obtained by permuting each neuron’s counts across transitions, which holds every neuron’s marginal rate fixed while abolishing cross-neuron coincidence within the real data (Methods). Claustral synchrony was elevated throughout the analysis window, consistent with correlated variability in the population, and rose to a transition-locked peak at the UP-to-DOWN transition (z = 7.59 versus a baseline of 4.69; P = 0.0004, n = 20 sessions, Wilcoxon signed-rank), precisely where the mean rate reached its minimum (Fig. 4b,c). While the population rate declined into the transition, the population fired increasingly in concert.

**Figure 4.**
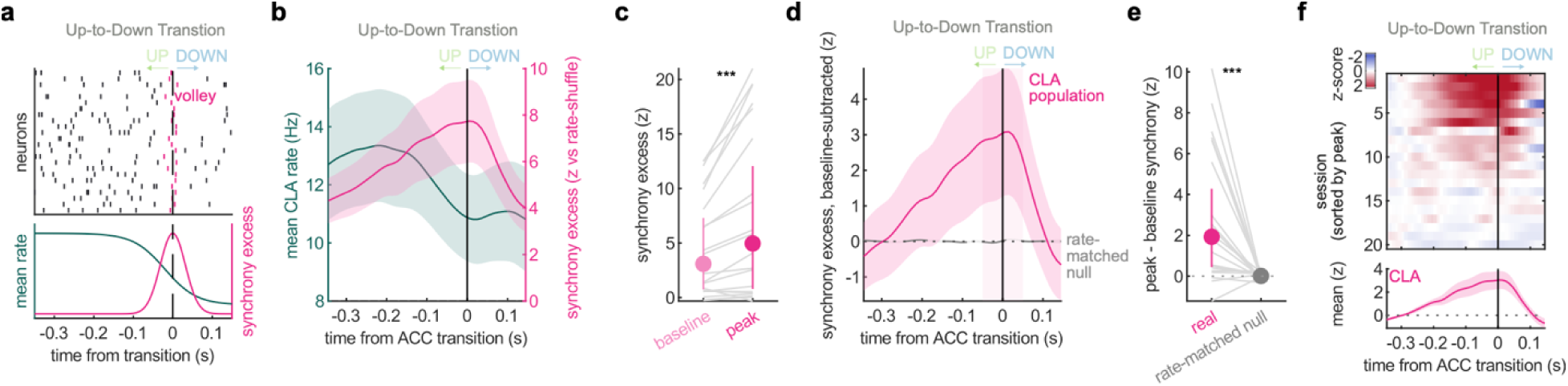
A coordinated claustral volley, not a rise in firing rate, accompanies the cortical UP-to-DOWN transition. (**a**) Schematic of the tested dissociation: a transition could be signaled either by an increase in the level of claustral firing or by a transient increase in its coordination, which are separable variables predicting different population signatures. (**b**) Population-mean claustrum firing rate (gold, left axis) and rate-controlled population synchrony excess (purple, right axis) aligned to the UP-to-DOWN transition. Synchrony excess is the cross-neuron spike coincidence per bin minus a rate-preserving shuffle that permutes each neuron’s counts across transitions, preserving marginal rate while abolishing coincidence, expressed as a z-score (Methods), and shown at its absolute level without baseline subtraction, so that the elevated resting excess is retained. Mean rate declines into the transition while synchrony excess peaks at it (peak within ±50 ms, z = 7.59, versus a baseline of 4.69 at −350 to −250 ms; Wilcoxon signed-rank P = 0.0004, n = 20 sessions). Shading, s.e.m. across sessions. (**c**) Synchrony excess at the transition peak (±50 ms) versus the pre-transition baseline (−350 to −250 ms) for each session, paired across sessions, the session-level form of the effect shown as an average in (b) (peak versus baseline, Wilcoxon signed-rank P = 0.0004, n = 20). Mean ± s.e.m. (**d**) Rate-matched specificity control. A synthetic population reproducing each neuron’s exact peri-transition firing-rate profile but spiking independently, with no imposed coordination, yields no synchrony excess. Baseline-subtracted synchrony excess over time, real claustrum population (purple) versus the rate-matched null (dashed grey), the latter remaining flat across the transition while the real population retains its peak. Shading, s.e.m. across sessions. (**e**) The same control summarized per session. Per-session peak-minus-baseline synchrony excess, real versus rate-matched null, paired by session (real 2.90 versus 0.01; Wilcoxon signed-rank P = 0.0006; null versus zero P = 0.79; n = 20). (**f**) Consistency across sessions. Synchrony excess for each session (rows, sorted by peak height), with each session referenced to its own pre-transition baseline (−350 to −250 ms) so that the transition-locked rise is shown against zero, in contrast to the absolute excess in (b); 17 of 20 sessions show a positive transition-locked peak. Population mean below; shading, s.e.m.

The permutation null removes coincidence at fixed rates but leaves open whether the specific shape of the peri-transition rate change could itself manufacture the peak. To rule this out, we constructed a synthetic population that reproduced each neuron’s exact peri-transition firing-rate profile but spiked independently, with no imposed coordination, and applied the identical measure. The synthetic population produced no synchrony excess, whereas the transition-locked excess of the real population substantially exceeded it (peak-minus-baseline 2.90 versus 0.01; paired P = 0.0006; synthetic versus zero P = 0.79; n = 20 sessions; Fig. 4d,e). A transition-locked rate modulation alone therefore cannot generate the effect, and the peak reflects genuine coordinated co-activation. This was consistent across recordings, with 17 of 20 sessions individually showing a positive transition-locked peak (Fig. 4f).

The coordinated event was specific to the UP-to-DOWN transition: at the DOWN-to-UP transition, synchrony rose together with the mean rate and showed no comparable dissociation (Extended Data Fig. 2). Entry into the DOWN state is thus marked by a selective coordination of claustral firing, paralleling the directional asymmetry of the claustral lead (Fig. 3c,d). Together, these results indicate that the claustral signal associated with the cortical UP-to-DOWN transition is carried through a transient coordination of firing that is invisible at the level of the mean rate: coordinated co-activation, not the firing-rate level, is the claustral variable most closely linked to the transition.

### Claustrum input directionally and causally controls cortical UP and DOWN state durations

These results identify the claustrum as a candidate source of the transition drive, carried by the coordinated volley characterized above, but do not establish that this input is required to produce the transition. To ask whether the CLA-to-ACC input is necessary, we photoinhibited ACC-projecting claustrum terminals during the peri-cue period (Fig. 5a) and asked, first, whether this abolished directed CLA-to-ACC coupling and, second, how it reshaped the cortical cycling.

**Figure 5.**
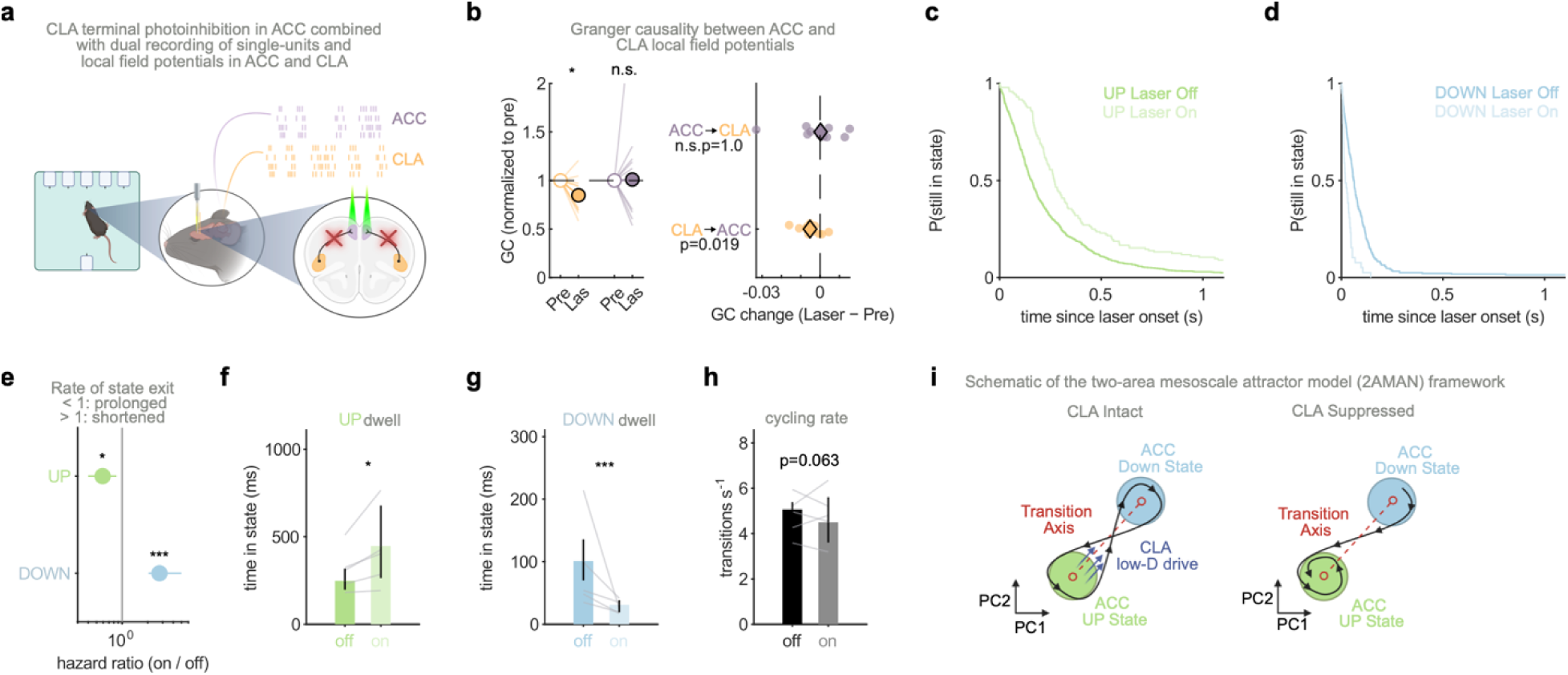
Claustrum input directionally and causally controls cortical state durations. (**a**) Schematic of experimental protocol. Bilateral CLA terminal photoinhibition in ACC during the peri-cue period(-0.5-1.5 s) of the cue-guided foraging task in mice. Photoinhibition was simultaneously combined with dual recording of single-units and local field potentials in ACC and CLA. (**b**) Granger causality between ACC and CLA local field potentials. **(Left)** Time-domain GC traces (ACC→CLA dark purple, CLA→ACC orange). CLA→ACC information flow was reduced under CLA inactivation (pre median = 0.0249, laser median = 0.0245; Wilcoxon signed-rank P = 0.019, n = 11 sessions), while ACC→CLA was unchanged (pre median = 0.0158, laser median = 0.0193; P = 1.0). **(Right)** Per-session GC changes (laser − pre). CLA→ACC shifts consistently below zero (median Δ = −0.0052, IQR [−0.007, −0.001]), while ACC→CLA scatters around zero (median Δ = 0.0003, IQR [−0.006, 0.004]). CLA inactivation selectively disrupts CLA→ACC information flow, consistent with CLA→ACC projections playing a causal role in coordinating ACC state dynamics. (**c**) Survival of the UP state from laser onset, conditioned on the state present at onset, for matched laser-on and laser-off epochs, plotted as the probability of remaining in the UP state against time from onset. The probability that an UP state survived the first 200 ms rose from 0.40 under laser-off to 0.69 under inhibition, indicating that cortex was held in the UP state longer when CLA was inactivated; the corresponding hazard ratio is given in (e). (**d**) Survival of the DOWN state from laser onset, plotted as in (c). The probability that a DOWN state survived the first 200 ms fell from 0.065 under laser-off to 0.000 under inhibition, indicating that cortex left the DOWN state sooner when CLA was inactivated; the corresponding hazard ratio is given in (e). (**e**) Exit hazard ratios for the UP and DOWN states under CLA inhibition (Cox proportional-hazards model, animal-clustered bootstrap 95% confidence intervals). The UP state was held longer (hazard ratio 0.58, interval 0.38 to 0.88, P = 0.011) and the DOWN state was exited sooner (hazard ratio 2.75, interval 2.04 to 4.98, P < 0.001); n = 144 laser-on and 669 laser-off UP entries and 39 laser-on and 278 laser-off DOWN entries, across 25 sessions and 10 animals. (**f**) Restricted mean time in state. The UP state lengthened from 248 to 447 ms under inhibition (+199 ms, animal-clustered bootstrap 95% interval 35 to 417 ms, P = 0.014). (**g**) Restricted mean time in state, as in (f). The DOWN state shortened from 101 to 31 ms (−70 ms, interval 40 to 109 ms, P < 0.001). (**h**) Per-window transition (cycling) rate, which fell modestly from 5.1 to 4.5 s⁻¹ under inhibition (roughly 11%, P = 0.063), consistent in direction with the dwell shifts in (f) and (**g**). Effects were unchanged when restricted to the cue-free interval preceding the chirp (Methods). (**i**) Schematic of the two-area mesoscale attractor model (2AMAN) framework (Recanatesi, 2022) illustrating the claustrum serving as a reciprocally connected subcortical structure with low-dimensional correlated variability with cortex. Claustral input to cortex is confined to a low-dimensional subspace aligned with the transition axis separating the cortical UP and DOWN attractors (left); this correlated drive moves the cortical state along that axis and promotes transitions between the two attractors. The cortical UP state is the network’s robust default attractor and the DOWN state is shallow and maintained by claustro-cortical drive; under claustral suppression (right) the low-dimensional drive is withdrawn, the UP attractor is stabilized and the DOWN attractor destabilized, so that the cortex dwells longer in UP and exits DOWN sooner, reproducing the photoinhibition phenotype. Quantification of the two-area model fit to the laser-off architecture and the single-parameter identification of claustro-cortical synaptic drive as the locus of the effect are shown in Extended Data Fig. 3.

Directed CLA-to-ACC coupling, a functional readout of the claustral drive to ACC, was quantified with time-domain Granger causality. Photoinhibition reduced CLA-to-ACC flow at the session level (signed-rank P = 0.019, n = 11 sessions) while leaving ACC-to-CLA flow unchanged (P = 1.0), with per-session changes tightly negative for CLA-to-ACC and zero-centered for ACC-to-CLA (Fig. 5b). The manipulation thus produced a directionally selective collapse of claustro-cortical coupling, consistent with the selective removal of CLA-to-ACC influence without altering cortical feedback to the claustrum.

Having confirmed that silencing removes the directed drive, we asked what that drive does to the cortical cycling, the time the network dwells in each state before switching. Locking to laser onset and conditioning on the state present at that moment, we measured the time to the first transition out of it, comparing laser-on with matched laser-off epochs (Methods). Suppression had opposite effects on the two states. When cortex was in the UP state at laser onset it remained there longer under inhibition: the hazard of leaving the UP state fell by roughly forty percent (Cox hazard ratio 0.58, animal-clustered bootstrap 95% confidence interval 0.38 to 0.88, P = 0.011; n = 144 laser-on and 669 laser-off UP entries across 25 sessions and 10 animals), and the probability that an UP state survived the first 200 ms of suppression rose from 0.40 to 0.69 (Fig. 5c,e). When cortex was in the DOWN state it left sooner: the hazard of exiting the DOWN state more than doubled (hazard ratio 2.75, bootstrap interval 2.04 to 4.98, P < 0.001; n = 39 and 278 DOWN entries; Fig. 5d,e). Removing the claustral drive therefore biased ACC away from the DOWN state in both phases of the cycle. These effects reflected spontaneous state holding rather than an altered cue response: both were unchanged when the analysis was restricted to the cue-free interval preceding the cue (data not shown). Because cue presentation falls within active foraging, moreover, the laser-off baseline here is the engaged regime characterized above (Fig. 2), so suppression prolongs UP states beyond their already elevated engaged duration.

Quantified as restricted mean time in state, suppression lengthened the UP state from 248 to 447 ms (an increase of 199 ms, animal-clustered bootstrap interval 35 to 417 ms, P = 0.014) and shortened the DOWN state from 101 to 31 ms (a decrease of 70 ms, interval 40 to 109 ms, P < 0.001; Fig. 5f,g). Because the UP lengthening exceeds the DOWN shortening, occupancy shifted toward the UP state and the cycle slowed modestly, the per-window transition rate falling by roughly 11% (5.1 to 4.5 per second; Fig. 5h), a change consistent in direction with the dwell shifts though at the margin of significance (P = 0.063).

These dwell-time effects have a parsimonious dynamical explanation. In a two-area mesoscale attractor model in which the claustrum supplies low-dimensional correlated drive aligned with the cortical UP-to-DOWN transition axis (Fig. 5i), fitting the model to the laser-off architecture and searching its parameters identified claustro-cortical synaptic drive as the only parameter whose reduction reproduced the photoinhibition phenotype on cortical cycling (Extended Data Fig. 3). The model thus recovers, from the geometry established in Fig. 3, the causal effect observed here. Together, these results establish ACC-projecting claustrum input as a directional drive that is necessary for timely entry into the cortical DOWN state and for limiting the duration of the UP state.

## Discussion

During flexible cue-guided behavior, state transitions within the metastable dynamics of frontal cortex appear to be a joint product of its reciprocal loop with the claustrum rather than a property of cortex alone. UP/DOWN cycling tracked behavioral engagement: cortex occupied a longer, more UP-dominated regime during active foraging than during disengagement, and the degree of UP-state dominance, indexed by the pre-cue UP fraction, predicted the probability of a correct cue-guided switch. Photoinhibition of ACC-projecting claustrum terminals selectively impaired cue-guided switching while sparing exploratory foraging. The claustrum led the UP-to-DOWN transition by ∼90 ms, delivered to ACC a drive confined to a low-dimensional subspace aligned with the cortical transition axis, and carried that drive not by an elevation of firing but by a transient rise in population synchrony as mean rate fell. Silencing this input reduced directed claustro-cortical coupling, biased cortex toward the UP state and away from the DOWN state, and modestly slowed the cycle. The claustrum thus supplies a directed drive that times entry into the cortical DOWN state, and this timing is required for efficient cue-guided switching.

These results identify the claustrum as a candidate biological implementation of the low-dimensional partner posited by mesoscale attractor models of cortical state^29^, in which a small, recurrence-poor area reciprocally coupled to cortex injects a correlated drive aligned with the axis separating successive attractors, generating transitions within the loop rather than imposing them on cortex from outside. The claustral drive to ACC met three signatures predicted for that partner: it was low-dimensional, it was aligned with the transition axis beyond chance, and it led rather than followed the transition. Fitting this model to the recorded cycling architecture, reducing the claustro-cortical drive was the only tested parameter whose reduction reproduced the silencing phenotype, identifying that drive as the locus of the effect (Extended Data Fig. 3) This role suits the claustrum’s architecture: its cortically projecting neurons are sparsely interconnected and lack electrical coupling, so the structure is unlikely to build attractors of its own and is instead positioned to read the cortex that drives it ^40^,while single neurons integrate convergent input from several frontal areas^32^.

The claustrum’s local circuitry offers a substrate for this operation. Cortico-claustral afferents recruit parvalbumin interneurons more strongly than the projection neurons they also contact, so cortical input arrives as excitation trailed closely by feedforward inhibition^40^,narrowing the window for effective summation and favoring correlated, near-simultaneous input over any single afferent ^41^. Claustral parvalbumin interneurons are themselves densely coupled by chemical and electrical synapses^40^ and could synchronize projection-neuron output, which targets the cortical parvalbumin and neuropeptide-Y interneurons best placed to promote a synchronous reset^33,34,42,43^. A claustrum organized this way — a coincidence detector whose output is coordination rather than rate — resolves an apparent contradiction with prior work: endogenous claustral firing fell as cortex approached the DOWN state, yet optogenetic claustral activation drives widespread cortical DOWN states^34^, If cortex responds to the coordination of claustral output rather than its level, optogenetic activation would promote the DOWN state by imposing synchrony directly, regardless of the accompanying change in rate. The operation linking ACC-projecting claustrum neurons to flexible behavior ^37,38^ may therefore be timing and coordination rather than overall activity level^44^.

More broadly, timing cortical state may be a general claustral operation, consistent with its proposed roles in perceptual binding, attention, and the coordination of cortical slow waves ^34,42,45,46^;our results extend this operation to state transitions in the waking, freely behaving animal. Within our task, the claustrum’s timing appears to maintain a behaviorally favorable regime: the engaged foraging regime lengthened and stabilized the UP state relative to disengagement, affording time for a cue to be read, and removing the claustral drive pushed the UP state past this optimum, delaying the reset that closes each processing epoch and impairing the switch^47^. In regulating the temporal boundaries of ACC activity states rather than the specific behavioral content they encode, the claustrum occupies much the role Crick and Koch anticipated when they cast it as a conductor coordinating otherwise independent cortical regions^46^, with a synchronized reset serving as the means by which one active pattern is cleared so that the next may prevail.

## Methods

### Animals

All procedures were conducted in accordance with the US National Institutes of Health Guide for the Care and Use of Laboratory Animals and were approved by the Institutional Animal Care and Use Committees at Oregon Health & Science University and the VA Portland Health Care System. A total of 19 wild-type C57BL/6J mice (10 female, 9 male; Jackson Laboratory) were used for all experiments. Animals were housed under standard laboratory conditions on a 12-hour reverse light–dark cycle with food available ad libitum. Water access was restricted to motivate task engagement as required.

### Surgical Procedures

All surgical procedures were performed in a single session including viral injection and chronic implant placement. Animals were anesthetized with vaporized isoflurane (3% induction, ∼ 1% maintenance) and head-fixed in a stereotaxic frame (David Kopf Instruments). Body temperature was maintained at 36–37 C using a feedback-controlled heating pad (Physitemp TCAT-2LV). Carprofen and dexamethasone were administered systemically at standard doses for analgesia and anti-inflammation; lidocaine was applied topically to the incision site prior to incision.

#### Dual-Virus Intersectional Strategy for Projection-Specific Inhibition

To selectively express inhibitory opsins in ACC-projecting claustrum neurons, a retrograde Cre-recombinase virus (AAVrg-hSyn-Cre-WPRE-hGH; Addgene 105553) was injected bilaterally into ACC, and a Cre-dependent ArchT virus (pAAV-FLEX-ArchT-tdTomato; Addgene 28305; AAV5) was injected bilaterally into the claustrum. This intersectional strategy restricts ArchT expression to claustrum neurons with a direct axonal projection to ACC. Control animals received an identical retrograde Cre injection into ACC, paired with a Cre-dependent fluorophore virus lacking an effector opsin (pAAV-FLEX-tdTomato; Addgene 28306; AAV5) injected into the claustrum.

ACC injections were made bilaterally at four sites (two per hemisphere) at the following coordinates relative to bregma: +1.9 mm anterior, ±0.4 mm mediolateral, at two depths ( −1.2 mm and −0.7 mm from brain surface), 200 nL per site. Claustrum injections were made bilaterally at one site per hemisphere (+ 1.45 mm anterior, ±2.65 mm mediolateral, −2.5 mm from brain surface, 50 nL per site). Viral incubation was allowed for a minimum of four weeks before recording to ensure sufficient expression.

#### Electrode Implant Design and Implantation

Custom 28-channel microwire implants were fabricated in-house. Electrode interface boards (EIB-36-Narrow-PTB; Neuralynx) were threaded with 12*μ*m diameter tungsten microwires (California Fine Wire Company) and secured with gold pins. Four additional larger-diameter wires were incorporated for LFP recordings from medial prefrontal cortex-connected regions (data not analyzed here). Silver wires were soldered to provide ground and reference signals. The assembly was mounted on a 3D-printed scaffold (Grey V4 resin; Formlabs) incorporating drivable adjustment screws allowing post-implant depth advancement. Of the 28 channels, 14 were targeted to ACC coordinates and 14 to claustrum coordinates. The ACC sub-bundle was co-housed with the ipsilateral optic fiber. Prior to implantation, all electrodes were gold-plated and impedance-matched to ∼200 kΩ at 1 kHz (nanoZ; White Matter LLC) to standardize yield. Implants with more than three open channels were discarded.

Electrode bundles were implanted in the left hemisphere at stereotaxic coordinates for ACC +1.9 mm anterior, ±0.4 mm mediolateral, −0.7 mm from brain surface) and claustrum ( +1.45 mm anterior, ±2.65 mm mediolateral, −2.5 mm from brain surface. A reference wire was implanted in the left striatum (+ 0.50 mm anterior, −1.60 mm lateral, −2.50 mm ventral relative to bregma), and a stainless-steel ground screw was placed over the contralateral cerebellum. All components were secured with dental cement (UNIFAST Trad; GC America) and skull-mounted support screws. Animals recovered for a minimum of one week before resuming water restriction and behavioral testing.

### Histological Verification and Inclusion Criteria

At the end of recording, electrolytic lesions were made through each electrode tip (10 *μ*A, 10 s, tip-positive) immediately before perfusion to mark recording locations. Brains were post-fixed in 4% paraformaldehyde, sectioned coronally at 50 *μ*m, and examined under fluorescence microscopy to confirm optic-fiber placement over ACC, viral (ArchT-tdTomato or tdTomato) expression in the claustrum, and electrode-bundle terminations. Claustrum boundaries were delineated with the Allen Mouse Brain Reference Atlas at the corresponding antero-posterior level; a lesion or expression locus was classified as claustral only if it fell entirely within the atlas-defined boundary, with loci in insular cortex, endopiriform nucleus, or striatum treated as off-target.

Animals or bundles failing histological verification were excluded from the corresponding analyses.

### Optogenetic Inhibition

Photoinhibition was delivered using a diode laser (Doric Lenses, 520 nm) appropriate for ArchT activation. Light was delivered continuously at 12 mW measured at the fiber tip. Bilateral optic fibers were implanted over ACC to achieve terminal photoinhibition of claustrum axons within ACC. To selectively target the cue-guided switching epoch, laser illumination was time-locked to the peri-cue window: onset 0.5 seconds before cue presentation, spanning the 1-second cue duration, and terminating 0.5 seconds after cue offset, for a total illumination duration of 2.0 seconds. Laser trials were interleaved pseudo-randomly with non-laser trials across both cue-present and cue-absent conditions. Control animals expressing only the fluorophore (tdTomato) underwent identical illumination protocols.

### Behavioral Apparatus and Task Design

Experiments were conducted in a custom-built six-port operant chamber (Sanworks Bpod State Machine r1.0; Sanworks LLC) equipped with illuminable, photo-gated, water-dispensing nose-poke ports. Task logic, event timing, port illumination, reward delivery, and synchronization with electrophysiological recordings were controlled by custom MATLAB scripts (MathWorks, Natick, MA). Auditory stimuli were generated by a Teensy microcontroller with a Teensy Audio Adaptor Board and delivered through a chamber-mounted speaker. TTL event codes from the Bpod microcontroller were co-recorded with neural data to enable precise alignment of behavioral events to neural signals.

#### Cued-switching Task

Animals were trained on a novel foraging with Cue-Guided switching task. Pseudo-random water reward was available at five front nose-poke ports, and an auditory cue (a 1-second linear frequency upsweep) signaled the transient availability of a substantially larger reward at a single back port (0.5 *μ*L front ports vs. 6 *μ*L back port). The pseudo-random front-port reward contingency followed a nearest-neighbor rule: upon reward delivery at one port, reward was reassigned to one of the two adjacent ports with equal probability, preventing stereotyped scanning while maintaining unpredictability. This contingency was adopted after pilot experiments revealed that fully random assignment induced default sequential scanning behavior.

Each session consisted of 100 trials. Trials began with a 10-second inter-trial interval (all ports off), after which all five front ports illuminated and foraging could commence. On cue-present trials (80% of all trials), the auditory cue was presented pseudo-randomly between 8 and 16 seconds after trial onset. Animals could respond by switching from foraging to the back port to retrieve reward within a 2.5-second window following cue offset. On cue-absent (forage-only) trials (20% of trials, randomly interleaved), no cue was presented and only front-port foraging was rewarded. Each trial terminated at t =35 seconds. Session length was fixed at 100 trials to prevent performance degradation due to satiation.

Prior to full task training, animals were pre-trained in stages: first learning the cue–back-port reward association via direct nose-poking, then the full cue-present/cue-absent contingency, and finally the complete task with active foraging. Animals were considered trained and eligible for testing when d exceeded 1.0 on at least three consecutive sessions, typically after 20–30 training sessions.

#### Trial Outcome Definitions

A trial was classified as a Hit if the animal visited the back port within the 2.5-second reward availability window following cue offset. A Miss was recorded if the animal failed to visit the back port within this window. A False Alarm was recorded if the animal visited the back port in the absence of a cue; such trials were terminated immediately and the next trial initiated. Trials on which the animal was at the back port immediately before cue onset were excluded from analysis to avoid contamination from carry-over position effects.

#### Behavioral Metrics

Hit rate was defined as the proportion of cue-present trials resulting in a rewarded back-port entry. False Alarm rate was defined as the rate of uncued back-port visits.

Discrimination performance (d’) was computed from hit rate *H* and false alarm rate *F* as 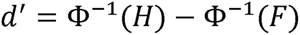, where Φ^-1^ denotes the inverse cumulative normal distribution. To avoid undefined values at extreme probabilities, rates were bounded away from 0 and 1 by a small constant prior to transformation. Front-port visit rate was computed as the number of front-port entries per unit time and used as a behavioral engagement covariate. A within-session laser effect metric (Δ*Hit* = *Hit*[*Laser*-*On*] – *Hit*[*Laser*-*Off*])was computed per session to control for session-level drift.

Continuous front-port poke rate was estimated for each trial using a sliding-window approach. Front-port entries were converted to an impulse train sampled at 2 kHz and convolved with a 2-second Hann window (area-normalized) to yield a smooth estimate of foraging rate in events s^-1^ .

### Behavioral Epoch Classification

To compare ACC dynamics across distinct behavioral contexts within a session, each peri-cue trial was segmented into foraging and non-foraging epochs based on the animal’s port-entry behavior. Foraging epochs were defined as periods during which the animal actively sampled front ports; non-foraging epochs were defined as periods during which the animal was not sampling any port for greater than 6 seconds. Epoch boundaries were determined from photogate-detected port entries and exits. For spectral analyses comparing foraging and non-foraging dynamics, equal-duration LFP segments were extracted from each epoch type within the same trial wherever possible to minimize cross-epoch confounds.

### Electrophysiology

Neural recordings were acquired using a CerePlex Direct acquisition system with a 32-channel CerePlex *μ* headstage (Blackrock Neurotech). Signals were sampled continuously at 30 kHz. Headstage cables and optic fiber tethers were routed via a Doric commutator to allow free movement. TTL pulses encoding behavioral event timestamps from the Bpod microcontroller were co-recorded on the same acquisition system to synchronize neural and behavioral data streams.

#### Spike Sorting and Single-Unit Inclusion

Single units were isolated offline using Kilosort3 and manually curated using Phy. Because exact electrode geometry within the microwire bundles was not precisely known, linear probe topology was assumed and drift-correction was disabled. Units automatically labeled “good” by Kilosort3 were reviewed and excluded if any of the following criteria were met: (1) waveform shape or amplitude drifted substantially across the recording session; (2) the unit was identified as a likely duplicate via cross-correlogram analysis; or (3) mean firing rate was below 0.5 Hz. Spike timestamps were exported to MATLAB for all subsequent analyses, organized by session, brain region, neuron identity, and trial. Neurons with insufficient variance across the analyzed epoch were excluded from population-level analyses.

### LFP Preprocessing

Raw broadband signals were high-pass filtered at 0.3 Hz and low-pass filtered at 500 Hz. LFP signals were downsampled to 2 kHz for all subsequent analysis. Band-limited signals used for UP/DOWN classification (below) and for directed-coupling analyses were obtained with zero-phase second-order Butterworth filters. Spectrograms were computed using a continuous wavelet transform with complex analytic wavelets (MATLAB’s cwt function) for display purposes.

### Spectral Parameterization of the ACC Field Potential

Periodic and aperiodic components of the ACC field-potential spectrum were separated with the specparam algorithm (formerly FOOOF; Donoghue et al., 2020). Power spectral densities (Welch) were exported from MATLAB to Python, fit with the specparam package, and the fitted components reimported for downstream statistics. Fits used a frequency range of 0.5–55 Hz, peak-width limits of 0.3–4.0 Hz, a maximum of four peaks, a minimum peak height of 0.05 (log power above the aperiodic fit), a peak threshold of 2.0 (relative SD of the smoothed flattened spectrum), and the “fixed” aperiodic mode (no knee). The model was fit once to the grand-mean spectrum for the headline aperiodic and peak parameters and once per session (SpectralGroupModel) for session-level statistics. Within each session the dominant oscillatory peak in each of four canonical bands (delta 0.5–4 Hz, theta 4–8 Hz, beta 8–30 Hz, gamma 30–55 Hz) was taken as the peak of largest aperiodic-corrected power whose center frequency fell within the band, yielding one peak per band per session; aperiodic-corrected peak power was compared across bands with a linear mixed-effects model (peak power band + (1 section)) with Bonferroni-corrected pairwise contrasts.

### UP/DOWN State Classification

Cortical UP and DOWN states were classified from the simultaneous LFP and multi-unit activity using a validated combined-evidence approach (Saleem et al, 2010). The 1–4 Hz analytic phase of the ACC LFP, computed via the Hilbert transform of a zero-phase bandpass-filtered signal, provided one source of evidence; the smoothed multi-unit activity (SMUA), obtained by summing spikes across all sorted ACC units within each 1 ms bin and convolving the result with a 25 ms Gaussian kernel, provided the second.

The two sources were combined into a single state-evidence variable *S_Comb_*(*t*) by averaging after standardization (Eq. 10 of Saleem et al, 2010):

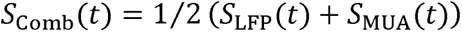

where *S*_LFP_(*t*) was derived from the analytic phase *ϕ*(*t*) of the bandpass-filtered LFP using the cosine model of Saleem et al, 2010 with the cross-validated phase offset 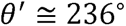 (Eq. 6), and *S*_MUA_(*t*) was the smoothed, [0,1]-normalized multi-unit signal. The phase offsets are fixed parameters imported from the published intracellular validation dataset and applied directly to the extracellular LFP; no intracellular recordings are required.

The combined evidence variable was fit with a two-component Gaussian mixture model (GMM) using the expectation-maximization algorithm, and each timepoint was assigned to UP or DOWN by maximum a posteriori probability. The lower-mean component was assigned to DOWN, the higher-mean component to UP. Per-sample labels were converted to state intervals without imposing a minimum dwell time, the duration distributions being unimodal and consistent with delta-band dynamics (median UP ≈ 0.23 s, median DOWN ≈ 0.09 s within foraging epochs); the brevity of the DOWN state reflects the asymmetric duty cycle of the delta band in awake, engaged cortex.

UP and DOWN dwell times were defined as the durations of continuously classified UP and DOWN epochs, and transition rate was defined as the number of state crossings per unit time within the analyzed window.

### UP/DOWN Architecture and Behavioral Engagement

The architecture of UP/DOWN cycling was compared between disengaged periods, when the animal was not sampling ports, and bouts of active foraging within the same laser-off trials (Hit and Miss pooled), using the behavioral epoch classification above and restricting to sessions with sufficient simultaneously recorded ACC units. For each epoch we computed the fraction of time spent in the UP state, UP and DOWN dwell times, and the transition rate, and compared foraging with disengagement using paired Wilcoxon signed-rank tests at the session level and Kolmogorov–Smirnov tests across the pooled distributions. To relate cortical state to behavior, the fraction of the pre-cue window spent in the UP state was entered as a predictor of trial outcome (correct versus incorrect) in a mixed-effects logistic regression with crossed random effects for session and trial, complemented by a Spearman rank correlation across trials.

### Transition Alignment and the Claustral Lead

Analyses were restricted to active-foraging epochs, defined as bouts of poke events separated by no more than 1 s and spanning at least 1 s, identically to the state-architecture analysis above. Each UP or DOWN transition whose departing state was contained within a foraging epoch was retained, and the aligned spike trains were masked to the interval spanning the departing-state onset and the arriving-state offset, clipped to the epoch boundary, so that no peri-transition estimate could include activity from a neighboring cortical cycle.

Per-neuron peri-transition histograms (UP-to-DOWN and DOWN-to-UP, computed separately) were constructed in 10 ms bins and z-scored across the supported window. For the population heatmap, the latency of maximal modulation was estimated from a random half of each neuron’s transitions and the displayed trace taken from the held-out half, a cross-validation that prevents the sort order from imposing spurious sequential structure; bins supported by fewer than five transitions were treated as missing.

The claustral lead was estimated as the lag maximizing the cross-correlation between the claustral and cortical population-mean traces, with positive values denoting a claustral lead, and was corroborated by the midpoint-crossing latencies of the two traces. Confidence intervals and one-sided *p*-values were obtained by a session-cluster bootstrap (5,000 resamples, sessions as the resampling unit). To remove the circularity arising from the cortical multi-unit contribution to state detection, a held-out procedure detected states from a random half of the ACC units and aligned the complementary half; a field-potential-only variant, which excluded multi-unit activity from detection entirely, was computed as an additional control (data not shown).

For the per-session lead (Fig 3d), the population-mean z-scored trace was formed from the claustrum and from the analyzed ACC units of each session, smoothed, and the modulation time taken as the time of maximal absolute slope within a window around the transition; claustrum and ACC times were compared by Wilcoxon signed-rank test (*n* = 13 sessions). Cortical transitions were detected from a held-out subset of ACC units and the analyzed ACC units were the complementary subset, so that the ACC modulation time was not fixed at the transition by construction.

### Low-Dimensional Claustro-Cortical Drive

The dimensionality and geometry of the claustrum-to-ACC drive were assessed in sessions with at least five simultaneously recorded units in each area. State-conditioned ACC population fluctuations were summarized by the participation ratio of the noise covariance, compared against a covariance-destroying shuffle of the same activity. The shared claustrum-to-ACC variability was estimated by reduced-rank regression of ACC on claustrum population activity within state epochs, with the cross-validated optimal rank defining a communication subspace. The cortical transition axis was defined as the direction in ACC population-activity space connecting the UP-and DOWN-state centroids, and its alignment with the communication subspace was compared against a random-subspace null, both in sample and under a split-half estimate that took the axis and the subspace from independent epochs.

### Synchrony at State Transitions

Population synchrony was quantified, in each peri-transition bin, as the mean number of coincident cross-neuron spike pairs per transition across the simultaneously recorded claustral ensemble, from which a rate-preserving null was subtracted: each neuron’s per-bin counts were permuted across transitions independently (200 permutations), preserving every neuron’s marginal rate while destroying cross-neuron coincidence, and the synchrony excess was expressed as a *z*-score relative to this null. Transition-locked modulation was tested by comparing the mean synchrony excess in a peri-transition window (within 50 ms of the transition) against a baseline window (−350 to −250 ms relative to the transition) across sessions (Wilcoxon signed-rank). Sessions with fewer than three simultaneously recorded claustral units were excluded from the synchrony analysis. As a specificity control, a synthetic population was constructed that reproduced each neuron’s peri-transition firing-rate profile but spiked independently, with no imposed coordination, and the identical measure was applied.

### Directed Coupling and State-Duration Survival

Time-domain Granger causality between ACC and CLA local field potentials was computed using bivariate vector autoregressive models on session-pooled, downsampled (200 Hz) LFP segments. Model order was selected per session by Bayesian information criterion. ACC→CLA and CLA→ACC log-likelihood ratios were computed in a pre-laser window and a matched laser window, and per-session changes were compared between conditions using paired Wilcoxon signed-rank tests.

To quantify how photoinhibition reshaped state durations, epochs were locked to laser onset and conditioned on the state present at that moment, and the time to the first transition out of that state was compared between laser-on and matched laser-off epochs. Exit hazards were estimated with a Cox proportional-hazards model with animal-clustered bootstrap confidence intervals, occupancy was summarized as the restricted mean time in each state, and the per-window transition rate was compared between conditions. Analyses were repeated within the cue-free interval preceding the chirp to separate spontaneous state holding from the cue-evoked response.

### Two-Area Attractor Model

We modelled ACC as a recurrent network of *N* = 1000 rate units with membrane currents *u_i_* obeying

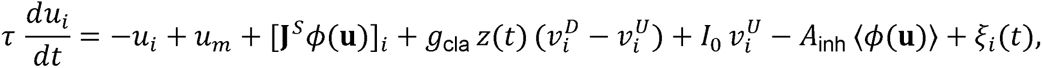

where the instantaneous rate is the single-neuron transfer function *ϕ* (*u_i_*) = 1/(1+exp(-*u_i –_ u_m)/_*β)). The midpoint *u_m_* =1.77 and inverse gain β=0.64 were obtained by least-squares fit of this logistic to the ACC current-to-rate transfer function inferred from the recorded UP and DOWN firing rates, so that the cortical input-output nonlinearity is data-derived rather than assumed (Recanatesi et al, 2022).

#### Attractor Storage

Two attractors, UP and DOWN, were defined by random binary patterns ξ*^u^* and ξ*^D^* with coding level *c* = 0.3 (each unit active independently with probability *c*); their centred forms are **v**^μ^ = ξ*^μ^*-*C*. Random patterns were used because the recorded UP and DOWN firing-rate vectors are strongly correlated (Pearson r ≈ 0.92) and would yield near-degenerate attractors, whereas the mesoscale framework stores effectively orthogonal patterns and fits the transfer function, not the patterns, to data. Patterns were stored Hebbian-style in a rank-two symmetric coupling,

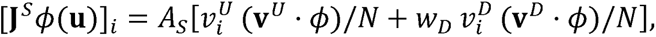

with retrieval gain A*_s_* = 28, set above the retrieval threshold 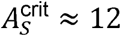 so that both patterns are stable fixed points in the absence of claustral drive. The DOWN storage weight *w_D_* = 0.3 (<1) renders DOWN a shallow, claustrum-maintained state rather than a co-equal attractor, encoding the hypothesis that the active cortical UP state is the default and that the claustrum promotes the silent DOWN state (Narikiyo et al, 2020); this asymmetry is what allows a reduction of claustral drive to lengthen UP and shorten DOWN jointly. A global inhibitory term 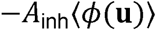 with *A_inh_* = 1.5 controlled total activity and enforced winner-take-all competition between patterns. An intrinsic UP-ignition term 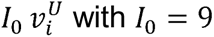 biased the cortex toward re-entry into UP and set the DOWN-to-UP transition, which in the model is claustrum-independent.

#### Claustral Drive

The claustrum was represented in reduced form by a single slow, low-dimensional drive *z*(*t*), the mesoscale correlated-variability term of the two-area model, acting through the fixed DOWN-promoting projection (**v**^D^ – **v**^U^) increasing *z* excites the DOWN pattern and suppresses the UP pattern, so that a coordinated claustral volley both entry into and maintenance of the cortical DOWN state. **z**(*t*) followed a rectified Ornstein-Uhlenbeck process,

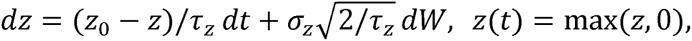

with baseline *z_0_ = 0* (attractors stable at rest), stationary standard deviation σ_z_ = 0.9, and timescale τ_z_ = 250 ms; the claustro-cortical drive gain was *g_cia_* = 10 in the control regime. The membrane time constant was τ = 12 ms and the per-unit private noise ξ*_i_*(*t*) was Gaussian with standard deviation 0.3 (in units of 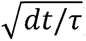). Transitions were thus noise-driven by the slow claustral volley, the attractors being stable when *z* was constant (Recanatesi et al, 2022).

#### Photoinhibition and Single-Parameter Search

Claustral photoinhibition was modelled as a reduction of the claustro-cortical drive, *g*_cia_ = 10 → 5.5, with all other parameters held at their control values; because the volley amplitude *σ_z_* and the gain *g*_cia_ both scale claustral output, an equivalent reduction of *σ_z_* produced identical effects. To test specificity, each parameter was altered once from the fixed control regime by a comparable fraction (claustro-cortical gain *g*_cia_ and volley amplitude *σ_z_* reduced 45%; attractor gain *A_S_* reduced to 18; global inhibition *A_inh_* reduced 50%; membrane time constant τ increased 60%; intrinsic ignition *I_0_* reduced 45%), and the resulting change in UP restricted mean survival time, DOWN restricted mean survival time, and cycling rate was computed relative to control. A manipulation was deemed to reproduce the laser effect if it lengthened UP, shortened DOWN, and slowed cycling simultaneously (Extended Data Figure [fig<u>:main_2aman]</u>).

#### State Segmentation and Numerical Implementation

Each model frame (10 ms, after downsampling the 0.5 ms integration) was labelled UP or DOWN by the larger of the two pattern overlaps 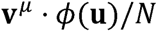. Dwell times were the durations of completed UP and DOWN epochs; survival functions were 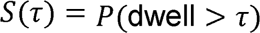 restricted mean survival times were the mean dwell with the right tail capped at 800 ms; exit-hazard ratios were the ratio of mean exit rates between the photoinhibition and control conditions for each state; and cycling rate was the number of completed UP-DOWN cycles per second. Equations were integrated by the Euler-Maruyama method with step *dt* =0.5 ms. Because the connectivity is rank-two by construction, the recurrent and claustral terms were evaluated as two length-*N* dot products per step rather than as an *N*x*N* matrix-vector product. Network connectivity was fixed by a single pattern seed and the dynamics by a single simulation seed; all reported quantities used 200 s of simulated time per condition (150 s for the parameter search). Simulations were written in Python with NumPy and SciPy and accelerated with Numba.

#### Model Scope

The model reproduces the direction of all three photoinhibition signatures and the magnitude of the UP effect (exit-hazard ratio 0.49 versus a recorded 0.58); the DOWN effect is weaker than observed (exit-hazard ratio 1.46 versus 2.75) because the network retains a finite minimum transition time and cannot abolish DOWN states as completely as photoinhibition does, and the fractional change in cycling exceeds the recorded value. The claustrum is represented as a reduced low-dimensional drive rather than as an explicit second-area population, and the gain and amplitude of that drive are not separately identifiable; the model therefore supports the conclusion that claustro-cortical synaptic drive is the parameter controlling the cortical cycling architecture, without resolving the cellular form of that drive.

### Statistical Analysis

Unless otherwise noted, statistical tests were nonparametric. Within-subject comparisons used two-tailed Wilcoxon signed-rank tests; between-group comparisons used two-tailed Wilcoxon rank-sum tests. Trial-level analyses with crossed random effects (session, trial, animal) were performed using generalized linear mixed-effects models implemented in MATLAB’s fitglme. Effect sizes are reported as median s.e.m. unless otherwise noted. The unit of analysis was the session for behavioral measures and the neuron for single-unit measures; exact sample sizes, test statistics, and *p*-values are reported in the Results and figure legends.

## End Notes

### Data Availability

The datasets generated during and analyzed during the current study are available from the corresponding author on reasonable request.

### Code Availability

All analyses and figure generation were performed using custom MATLAB code (MathWorks, Natick, MA). The code generated for the current study are available from the corresponding author on reasonable request.

### Author Contributions

(conceptualization, methodology, investigation, formal analysis, writing, supervision, funding acquisition) map onto it naturally if you want a checklist to work from.

R.J.O, and A.I.A. conceptualized and designed the experiments; R.J.O performed the experiments; R.J.O, and L.M. analyzed the data; R.J.O. wrote the initial draft of the manuscript; R.J.O, A.S., L.B., R.M. and A.I.A. revised the manuscript and approved the final version before submission.

## References

1. Mazzucato, L., Fontanini, A. & La Camera, G. Dynamics of Multistable States during Ongoing and Evoked Cortical Activity. J. Neurosci. 35, 8214–8231 (2015).

2. La Camera, G., Fontanini, A. & Mazzucato, L. Cortical computations via metastable activity. Curr. Opin. Neurobiol. 58, 37–45 (2019).

3. Finkelstein, A. et al. Attractor dynamics gate cortical information flow during decision-making. Nat. Neurosci. 24, 843–850 (2021).

4. Steriade, M., Nuñez, A. & Amzica, F. A novel slow (&< 1 Hz) oscillation of neocortical neurons in vivo: depolarizing and hyperpolarizing components. J. Neurosci. Off. J. Soc. Neurosci. 13, 3252–3265 (1993).

5. Sanchez-Vives, M. V. & McCormick, D. A. Cellular and network mechanisms of rhythmic recurrent activity in neocortex. Nat. Neurosci. 3, 1027–1034 (2000).

6. Petersen, C. C. H., Hahn, T. T. G., Mehta, M., Grinvald, A. & Sakmann, B. Interaction of sensory responses with spontaneous depolarization in layer 2/3 barrel cortex. Proc. Natl. Acad. Sci. 100, 13638–13643 (2003).

7. Poulet, J. F. A. & Petersen, C. C. H. Internal brain state regulates membrane potential synchrony in barrel cortex of behaving mice. Nature 454, 881–885 (2008).

8. Vyazovskiy, V. V. et al. Local sleep in awake rats. Nature 472, 443–447 (2011).

9. Harris, K. D. & Thiele, A. Cortical state and attention. Nat. Rev. Neurosci. 12, 509– 523 (2011).

10. McGinley, M. J. et al. Waking State: Rapid Variations Modulate Neural and Behavioral Responses. Neuron 87, 1143–1161 (2015).

11. Cascella, N. G., Gerner, G. J., Fieldstone, S. C., Sawa, A. & Schretlen, D. J. The insula-claustrum region and delusions in schizophrenia. Schizophr. Res. 133, 77–81 (2011).

12. Bernstein, H.-G. et al. Bilaterally reduced claustral volumes in schizophrenia and major depressive disorder: a morphometric postmortem study. Eur. Arch. Psychiatry Clin. Neurosci. 266, 25–33 (2016).

13. Kang, S. S. & Sponheim, S. 55. Abnormal Claustral–Cortical Functional Brain Networks in People With Schizophrenia During Perception and Resting: Brain Network Dynamics Underlying Psychotic Symptoms. Schizophr. Bull. 43, S28–S29 (2017).

14. Arieli, A., Sterkin, A., Grinvald, A. & Aertsen, A. Dynamics of ongoing activity: explanation of the large variability in evoked cortical responses. Science 273, 1868– 1871 (1996).

15. Poulet, J. F. A. & Crochet, S. The Cortical States of Wakefulness. Front. Syst. Neurosci. 12, 64 (2019).

16. McGinley, M. J., David, S. V. & McCormick, D. A. Cortical Membrane Potential Signature of Optimal States for Sensory Signal Detection. Neuron 87, 179–192 (2015).

17. Jercog, D. et al. UP-DOWN cortical dynamics reflect state transitions in a bistable network. eLife 6, e22425 (2017).

18. Neske, G. T. The Slow Oscillation in Cortical and Thalamic Networks: Mechanisms and Functions. Front. Neural Circuits 9, 88 (2015).

19. Friston, K. J. Transients, Metastability, and Neuronal Dynamics. NeuroImage 5, 164–171 (1997).

20. Rushworth, M. F. S., Walton, M. E., Kennerley, S. W. & Bannerman, D. M. Action sets and decisions in the medial frontal cortex. Trends Cogn. Sci. 8, 410–417 (2004).

21. Kennerley, S. W., Walton, M. E., Behrens, T. E. J., Buckley, M. J. & Rushworth, M. F. S. Optimal decision making and the anterior cingulate cortex. Nat. Neurosci. 9, 940–947 (2006).

22. Tervo, D. G. R. et al. The anterior cingulate cortex directs exploration of alternative strategies. Neuron 109, 1876–1887.e6 (2021).

23. Balleine, B. W. & Dickinson, A. Goal-directed instrumental action: contingency and incentive learning and their cortical substrates. Neuropharmacology 37, 407–419 (1998).

24. Proskurin, M., Manakov, M. & Karpova, A. ACC neural ensemble dynamics are structured by strategy prevalence. eLife 12, e84897 (2023).

25. Alizadeh Mansouri, F., Buckley, M. J. & Tanaka, K. Mapping causal links between prefrontal cortical regions and intra-individual behavioral variability. Nat. Commun. 15, 140 (2024).

26. Durstewitz, D., Vittoz, N. M., Floresco, S. B. & Seamans, J. K. Abrupt Transitions between Prefrontal Neural Ensemble States Accompany Behavioral Transitions during Rule Learning. Neuron 66, 438–448 (2010).

27. Hasenstaub, A., Sachdev, R. N. S. & McCormick, D. A. State Changes Rapidly Modulate Cortical Neuronal Responsiveness. J. Neurosci. 27, 9607–9622 (2007).

28. Compte, A., Sanchez-Vives, M. V., McCormick, D. A. & Wang, X.-J. Cellular and network mechanisms of slow oscillatory activity (&<1 Hz) and wave propagations in a cortical network model. J. Neurophysiol. 89, 2707–2725 (2003).

29. Recanatesi, S., Pereira-Obilinovic, U., Murakami, M., Mainen, Z. & Mazzucato, L. Metastable attractors explain the variable timing of stable behavioral action sequences. Neuron 110, 139–153.e9 (2022).

30. Jackson, J., Smith, J. B. & Lee, A. K. The Anatomy and Physiology of Claustrum-Cortex Interactions. Annu. Rev. Neurosci. 43, 231–247 (2020).

31. Wang, Q. et al. Regional and cell-type-specific afferent and efferent projections of the mouse claustrum. Cell Rep. 42, 112118 (2023).

32. Shelton, A. M. et al. Single neurons and networks in the claustrum integrate input from widespread cortical sources. 2022.05.06.490864 Preprint at 10.1101/2022.05.06.490864 (2022).

33. Jackson, J., Karnani, M. M., Zemelman, B. V., Burdakov, D. & Lee, A. K. Inhibitory Control of Prefrontal Cortex by the Claustrum. Neuron 99, 1029–1039.e4 (2018).

34. Narikiyo, K. et al. The claustrum coordinates cortical slow-wave activity. Nat. Neurosci. 23, 741–753 (2020).

35. Atlan, G. et al. Claustrum neurons projecting to the anterior cingulate restrict engagement during sleep and behavior. Nat. Commun. 15, 5415 (2024).

36. Atlan, G. et al. The Claustrum Supports Resilience to Distraction. Curr. Biol. CB 28, 2752–2762.e7 (2018).

37. Peretz-Rivlin, N. et al. Claustral Input Consolidates Anterior Cingulate Cortical States for Selective Behavior. 2026.01.07.698269 Preprint at 10.64898/2026.01.07.698269 (2026).

38. White, M. G. et al. Anterior Cingulate Cortex Input to the Claustrum Is Required for Top-Down Action Control. Cell Rep. 22, 84–95 (2018).

39. Saleem, A. B., Chadderton, P., Apergis-Schoute, J., Harris, K. D. & Schultz, S. R. Methods for predicting cortical UP and DOWN states from the phase of deep layer local field potentials. J. Comput. Neurosci. 29, 49–62 (2010).

40. Kim, J., Matney, C. J., Roth, R. H. & Brown, S. P. Synaptic Organization of the Neuronal Circuits of the Claustrum. J. Neurosci. 36, 773–784 (2016).

41. Bruno, R. M. Synchrony in sensation. Curr. Opin. Neurobiol. 21, 701–708 (2011).

42. Do, A. D., Portet, C., Goutagny, R. & Jackson, J. The claustrum and synchronized brain states. Trends Neurosci. 47, 1028–1040 (2024).

43. Cho, K. K. A., Shi, J., Phensy, A. J., Turner, M. L. & Sohal, V. S. Long-range inhibition synchronizes and updates prefrontal task activity. Nature 617, 548–554 (2023).

44. Madden, M. B. et al. A role for the claustrum in cognitive control. Trends Cogn. Sci. 26, 1133–1152 (2022).

45. Goll, Y., Atlan, G. & Citri, A. Attention: the claustrum. Trends Neurosci. 38, 486–495 (2015).

46. Crick, F. C. & Koch, C. What is the function of the claustrum? Philos. Trans. R. Soc. B Biol. Sci. 360, 1271–1279 (2005).

47. Buzsáki, G. Neural syntax: cell assemblies, synapsembles, and readers. Neuron 68, 362–385 (2010).

