## Supplemental Figures and Legends for "A claustro-cortical loop times state transitions for flexible behavior"

**Extended Data**

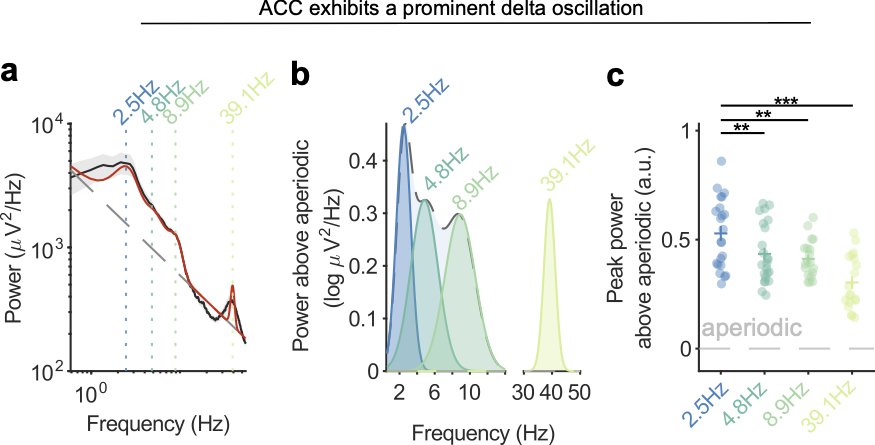

**Extended Data Figure 1. ACC exhibits a prominent delta oscillation.**

(**a**) Grand-mean ACC power spectral density (Welch, log–log; mean ± s.e.m. across sessions) overlaid with the FOOOF aperiodic component (dashed grey) and full model fit (red); aperiodic exponent = 0.69, offset = 3.46, R² = 0.969 (n = 25 sessions). Vertical lines mark identified oscillatory peaks at 2.5, 4.8, 8.9 and 39.1 Hz.

(**b**) Aperiodic-corrected oscillatory component (FOOOF peaks-only fit) showing the four peaks identified in (a) (delta CF = 2.46 Hz, BW = 1.64 Hz; theta CF = 4.83 Hz, BW = 3.09 Hz; beta CF = 8.88 Hz, BW = 3.45 Hz; gamma CF = 39.15 Hz, BW = 4.00 Hz).

(**c**) Per-session FOOOF peak power above aperiodic for each band; delta exhibited significantly greater peak power than theta, beta and gamma (linear mixed-effects model, peak power ~ band + (1 | session), Bonferroni-corrected: delta versus theta P = 0.003, delta versus beta P = 0.001, delta versus gamma P < 0.001; n = 25 sessions).

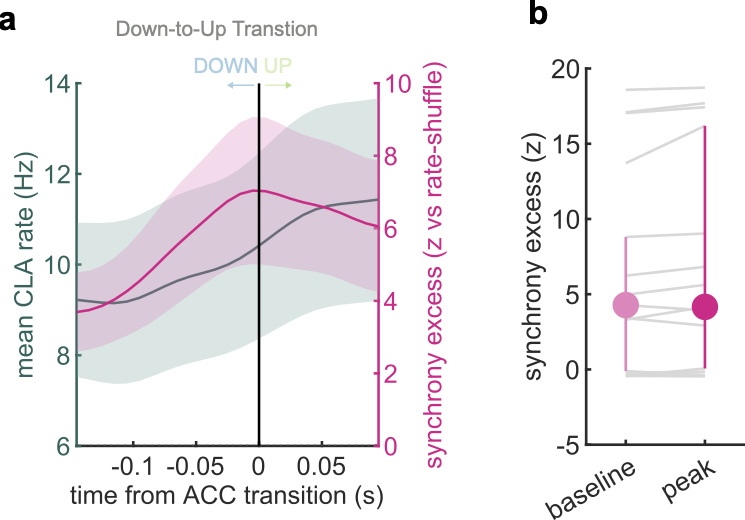

**Extended Data Figure 2. Claustral synchrony does not dissociate from firing rate at the DOWN-to-UP transition.**

Companion analysis to Fig. 4, applying the identical rate-controlled synchrony measure (Methods) to the cortical DOWN-to-UP transition, the transition at which the claustrum shows no resolvable lead (Fig. 3c,d).

(**a**) Population-mean claustrum firing rate (teal, left axis) and rate-controlled population synchrony excess (z versus a rate-preserving shuffle; magenta, right axis) aligned to the ACC DOWN-to-UP transition (time zero). In contrast to the UP-to-DOWN transition (Fig. 4b), where synchrony peaks as the mean rate falls, here mean rate and synchrony rise together across the transition, with no dissociation between them. Shading, s.e.m. across sessions (n = 20).

(**b**) Synchrony excess at the pre-transition baseline (−350 to −250 ms) versus the transition peak (±50 ms) for each session (grey lines), with the session summary (mean ± s.e.m.). No selective transition-locked coordination beyond the accompanying rise in rate is present, so the coordinated volley identified in Fig. 4 is specific to the UP-to-DOWN transition (n = 20 sessions).

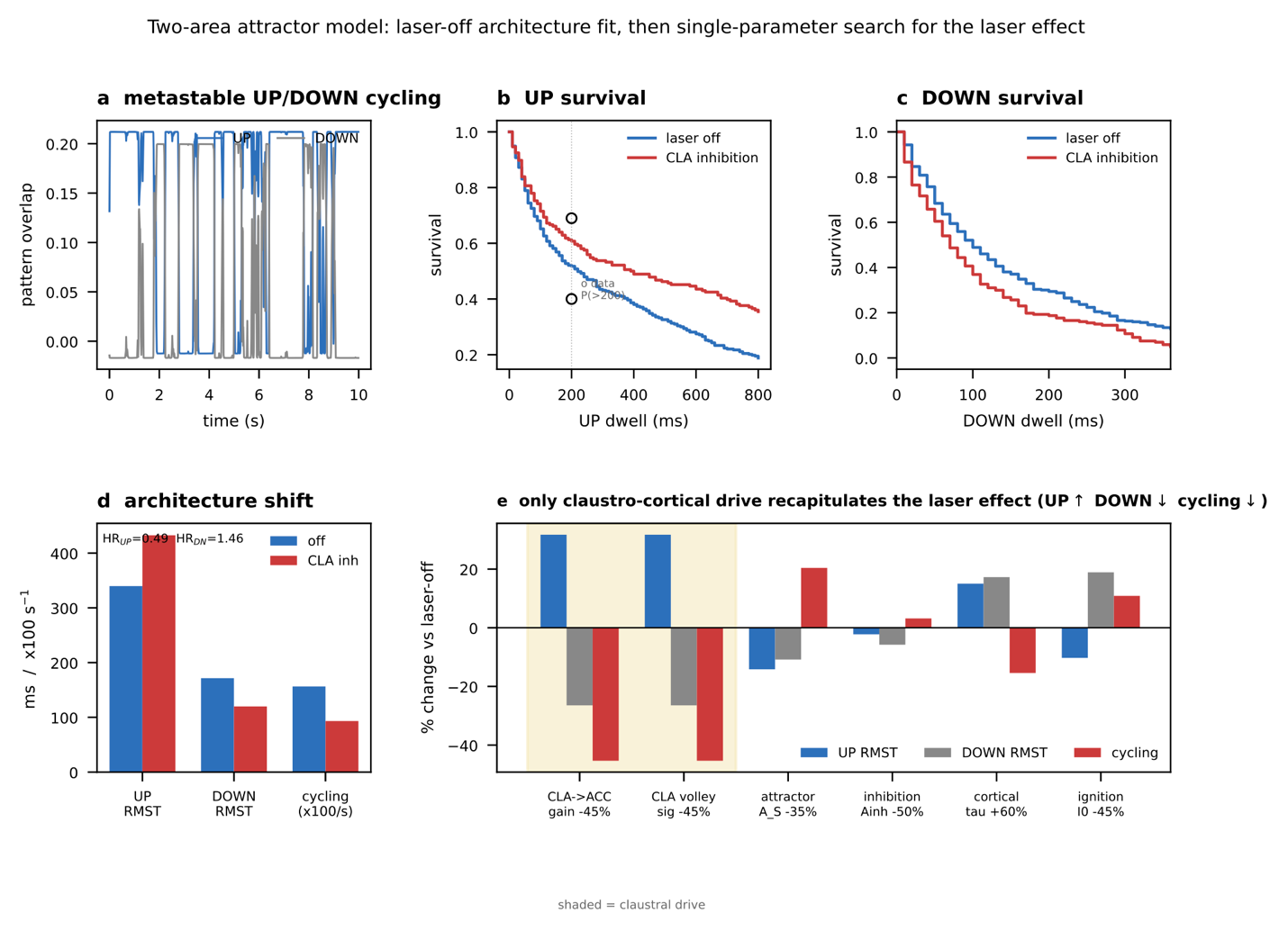

**Extended Data Figure 3. A two-area attractor model identifies claustro-cortical synaptic drive as the locus of the photoinhibition effect.**

Quantification of the two-area mesoscale attractor model (2AMAN) schematized in Fig. 5i, in which ACC is a recurrent network storing UP and DOWN attractors and the claustrum supplies a slow, low-dimensional drive aligned with the transition axis (Recanatesi et al., 2022; Methods). The model was fit to the laser-off (control) architecture, and photoinhibition was modelled as a reduction of the claustro-cortical drive (g_cla, or the equivalent volley amplitude σ_z), with all other parameters held at their control values.

(**a**) Example of metastable UP/DOWN cycling in the fitted laser-off regime (pattern overlap over time). The control fit reproduced the recorded architecture without reference to the photoinhibition data: UP fraction 0.73 (recorded 0.73), right-skewed dwell-time distributions, and median UP and DOWN durations of 220 and 100 ms, bracketing the recorded restricted mean survival times of 248 and 101 ms.

(**b**) UP-state survival function for the control (laser-off) and reduced-drive (CLA inhibition) regimes; open symbols, recorded data; P(>200), probability of surviving the first 200 ms. Reducing the claustro-cortical drive raised UP survival (UP exit-hazard ratio HR_UP = 0.49, closely matching the recorded 0.58; recorded 95% CI 0.38 to 0.88).

(**c**) DOWN-state survival function, plotted as in (b). Reducing the drive lowered DOWN survival, moving the DOWN exit-hazard ratio in the observed direction (HR_DN = 1.46; recorded 2.75); the model effect is weaker than observed because the network retains a finite minimum transition time.

(**d**) Architecture shift produced by reducing the claustro-cortical drive: UP restricted mean survival time lengthened, DOWN shortened, and cycling slowed, jointly reproducing the three signatures of the photoinhibition phenotype (HR_UP = 0.49, HR_DN = 1.46).

(**e**) Single-parameter specificity search. Percent change from the laser-off regime in UP restricted mean survival time (RMST), DOWN RMST, and cycling rate when each parameter was altered once by a comparable amount (claustro-cortical gain g_cla and volley amplitude σ_z reduced 45%; attractor gain A_S reduced 35%; global inhibition A_inh reduced 50%; membrane time constant τ increased 60%; intrinsic ignition I_0 reduced 45%). Only reduction of the claustro-cortical drive (shaded) lengthened UP, shortened DOWN, and slowed cycling simultaneously (UP RMST +32%, DOWN −26%, cycling −45%); no intrinsic-cortical parameter reproduced the phenotype. The photoinhibition effect therefore maps onto a reduction of claustro-cortical synaptic drive and onto no other parameter in the model.
